# Global Genome Nucleotide Excision Repair is Organised into Domains Promoting Efficient DNA Repair in Chromatin

**DOI:** 10.1101/050807

**Authors:** Shirong Yu, Katie Evans, Patrick van Eijk, Mark Bennett, Richard M. Webster, Matthew Leadbitter, Yumin Teng, Raymond Waters, Stephen P. Jackson, Simon H. Reed

**Affiliations:** Present address: Cambridge Epigenetix, Jonas Webb Building, Babraham Campus, Cambridge, CB22 3AT; Present address: Crescendo Biologics Ltd, Meditrina Building 260, Babraham Research Campus, Cambridge, CB22 3AT; Wellcome Trust/Cancer Research UK Gurdon Institute, The Henry Wellcome Building of Cancer and Developmental Biology, University of Cambridge, Tennis Court Road, Cambridge, CB2 1QN

**Keywords:** Global Genome Nucleotide Excision Repair, UV, genome structure, chromatin remodelling, yeast

## Abstract

The rates at which lesions are removed by DNA repair can vary widely throughout the genome with important implications for genomic stability. To study this, we measured the distribution of nucleotide excision repair (NER) rates for UV-induced lesions throughout the budding yeast genome. By plotting these repair rates in relation to genes and their associated flanking sequences, we reveal that in normal cells, genomic repair rates display a distinctive pattern, suggesting that DNA repair is highly organised within the genome. Furthermore, by comparing genome-wide DNA repair rates in wild-type cells, and cells defective in the global genome-NER (GG-NER) sub-pathway, we establish how this alters the distribution of NER rates throughout the genome. We also examined the genomic locations of GG-NER factor binding to chromatin before and after UV irradiation revealing that GG-NER is organised and initiated from specific genomic locations. At these sites, chromatin occupancy of the histone acetyl transferase Gcn5 is controlled by the GG-NER complex, which regulates histone H3 acetylation and chromatin structure, thereby promoting efficient DNA repair of UV-induced lesions. Chromatin remodeling during the GG-NER process is therefore organized into these genomic domains. Importantly, loss of Gcn5, significantly alters the genomic distribution of NER rates, a finding that has important implications for the effects of chromatin modifiers on the distribution of mutations that arise throughout the genome.

## Introduction

DNA, the key molecule of heredity, is susceptible to damage to its structure because it is continually exposed to the deleterious effects of normal cellular metabolic processes and external genotoxic stresses, such as ultraviolet (UV) radiation and chemical damage (Friedberg 2003). Thousands of lesions occur every day in the DNA of each of our cells, the immediate implications of which include disruption of DNA replication and cell division as well as defective gene regulation. Long-term effects include the introduction of DNA mutations, which alter the genetic information of the cell. Repair of damaged DNA is therefore fundamental to the maintenance of genome stability (Holmquist and Gao 1997). Whole-exome sequencing studies of the range of human cancer types (The Cancer Genome Atlas Research et al. 2013) identified tumour-specific somatic mutations and multiple mutational signatures associated with the different cancer types (Alexandrov et al. 2013a). The causes of these mutational signatures in tumours falls broadly into two groups that arise from exposure of cells to environmental mutagens, such as UV light or polycyclic aromatic hydrocarbons from cigarette smoke; or from defects in the various DNA repair pathways (Nik-Zainal et al. 2012; Alexandrov et al. 2013a; Alexandrov et al. 2013b). Collectively, these observations demonstrate the importance of understanding how genome damage is formed and efficiently and accurately repaired in normal cells.

Nucleotide excision repair (NER) acts on a spectrum of various types of DNA damage that have the common property of distorting the normal features of the DNA double-helix. Over thirty polypeptides are involved in the basic NER reaction processes. Two damage-recognition pathways operate: the rapid acting transcription coupled repair pathway (TC-NER) that operates on the transcribed strands of actively transcribing genes and involves RNA polymerase II in the damage recognition step; and the slower acting global genome repair pathway (GG-NER) that operates on all DNA, including non-transcribed and repressed regions, and involves a subset of proteins in the early stages of DNA damage recognition (Fousteri and Mullenders 2008). Following the initial stages of DNA damage detection, these two pathways converge and utilise the same set of DNA repair proteins. The majority of yeast NER genes have well conserved structural and/or functional human homologues, and the main features of both GG-NER and TC-NER pathways are evolutionary conserved (Hoeijmakers 1993; Hoeijmakers 1994).

In the nucleus, DNA is packaged into the nucleoprotein complex of chromatin. At present, how NER operates on naked DNA is quite well understood, but our knowledge of how the process operates in cellular chromatin is still emerging (Adam et al. 2015). Determining how DNA damage is sensed and removed from DNA packaged into chromatin is central to advancing our understanding of the mechanisms that promote genome stability and their effect on human health. Recent advances are providing important insights into such responses (Adam et al. 2015; Polo 2015) and we previously identified a complex of the *Saccharomyces cerevisiae* proteins Rad7, Rad16 and Abf1, that is required specifically for GG-NER in yeast. We demonstrated a novel function of the Abf1 component of this complex in NER, and showed that efficient GG-NER requires Abf1 to be bound to specific consensus DNA binding sites (Reed et al. 1999), which can be found at hundreds of locations throughout the yeast genome (Yu et al. 2009). The Rad16 protein is a member of the SWI/SNF super-family of chromatin remodelling factors. Proteins in this super-family contain conserved ATPase motifs and have been identified as subunits of protein complexes that demonstrate chromatin-remodelling activity (Flaus and Owen-Hughes 2011). Since Rad16 operates on repressed and non-transcribed regions of the genome in GG-NER, it has long been assumed that its role might involve chromatin remodelling (Verhage et al. 1994), which conceivably promotes improved accessibility of damaged DNA within chromatin to promote efficient DNA repair. Rad16 also contains a C3HC4 type RING domain, which is an important functional domain in ubiquitin E3 ligase proteins. The Rad7 protein also has two structural motifs: a leucine-rich repeat domain involved in protein-protein interactions and a SOCS-box domain that comprises another ubiquitin E3 ligase motif. Indeed, we have previously reported that the GG-NER complex has *bona fide* E3 ubiquitin ligase activity involving the Cul3 and Elc1 proteins (Pintard et al. 2004; Willems et al. 2004; Gillette et al. 2006).

Previously, we investigated how the yeast GG-NER complex remodels chromatin by examining events at a single genetic locus (Yu et al. 2011). This work has established that the complex promotes UV-induced chromatin remodelling necessary for DNA repair by recruiting the histone acetyl transferase (HAT) Gcn5 onto chromatin, which promotes increased histone H3 acetylation levels that in turn alter chromatin structure (Yu et al. 2011). These observations offered important insights into how the GG-NER complex promotes the UV-induced chromatin remodelling necessary for DNA repair at the genetic locus examined.

In the present study we carry out an expanded investigation of these parameters to examine how the GG-NER process is organised throughout the entire yeast genome. To tackle this issue we developed a genome-wide DNA repair assay based on ChIP-Chip, referred to as 3D-DIP-Chip (Teng et al. 2011; Powell et al. 2015). The method relies upon immuno-affinity capture of UV-induced DNA damage and the separation of damaged from undamaged DNA. The genomic locations of the DNA damage can then be identified by hybridisation of fluorescently labelled DNA to whole-genome DNA microarrays. These microarrays are optically scanned where fluorescence intensities are converted to numerical values that reveal a map of both the levels and locations of DNA damage over the genome. Repeating this process at different times after the induction of DNA damage permits an estimate of the relative rates of DNA repair to be made at individual sites throughout the genome (Teng et al. 2011; Powell et al. 2015). Comparison of variations in location specific DNA repair rates in wild-type and various mutant strains can then be made. We also measured the chromatin binding of the individual GG-NER factors, HAT occupancy, and histone H3 acetylation levels in chromatin throughout the genome, before and after UV irradiation, to understand how these events are organised.

Our results demonstrate that chromatin remodelling and repair is initiated from ABF1 DNA binding sites, controlled by the components of the GG-NER complex found predominantly at many hundreds of intergenic regions throughout the genome. We reveal that the Rad7-Rad16-Abf1 GG-NER complex is chromatin-bound at ABF1 binding sites in the absence of UV damage, suggesting that the co-localisation of the components at these locations primes the genome for repair. Following UV irradiation, we show a UV-induced relocation of Rad7 and Rad16 that controls the level and distribution of the HAT Gcn5 on the chromatin in the vicinity of Abf1 binding sites. In this way, UV-induced acetylation of histone H3 is organised into domains around Abf1 binding sites, thus regulating chromatin structure in response to DNA damage. We show that this series of Rad7 and Rad16 dependent events is required to ensure the proper organisation of the GG-NER process throughout the genome. We discuss how these observations could help to explain how driver mutations in novel cancer genes involved in regulating chromatin structure contribute to altered patterns of genomic instability throughout the genome during tumourigenesis.

## Results

### The GG-NER complex promotes efficient repair of UV-induced DNA damage in non-transcribed genomic regions

The GG-NER pathway operates on all DNA, including non-transcribed and repressed regions of the genome. Here, we used 3D-DIP-Chip (Teng et al. 2011; Powell et al. 2015) to measure UV-induced DNA damage throughout the yeast genome at various time points after UV irradiation, to investigate the role of GG-NER in promoting removal of this damage. We previously developed the R software package Sandcastle (Bennett et al. 2015) for the analysis of this type of data, and it has been used to create the plots described here. As described previously (Teng et al. 2011), we observe a heterogeneous distribution of CPDs throughout the genome immediately after UV irradiation (Figure 1A). To calculate relative rates of CPD removal throughout the genome, we repeated the 3D-DIP-Chip procedure with DNA from cells which had been allowed two hours repair time after UV-irradiation and then subtracted these values from CPD levels immediately after UV irradiation to generate a genome-wide pattern of repair. As shown previously, we also observe a heterogeneous distribution of DNA repair rates for the removal of CPDs in relation to the linear arrangement of the genome (Figure 1B) (Teng et al. 2011; Powell et al. 2015). Therefore, to examine the distribution of repair rates in relation to gene structure, we produced composite gene open-reading frame (ORF) plots including their flanking regions to derive an average trend of the data (Described in Figure 1C). Thus ORFs ranging from 500 to 1500 bp were used to generate composite plots of DNA repair in this context, including DNA sequences ranging up to 2 kbp upstream and downstream of the ORFs. This represents approximately 85% coverage of the yeast genome. It is important to note that the profile-plotting function in Sandcastle ensures that no region of the genome is represented more than once in these plots. This feature is important in preventing the duplication of genomic data where the regions plotted overlap (as illustrated in Figure 1C). We refer to this style of figure as a ‘composite plot’ throughout the remainder of the manuscript.

**Figure 1.**
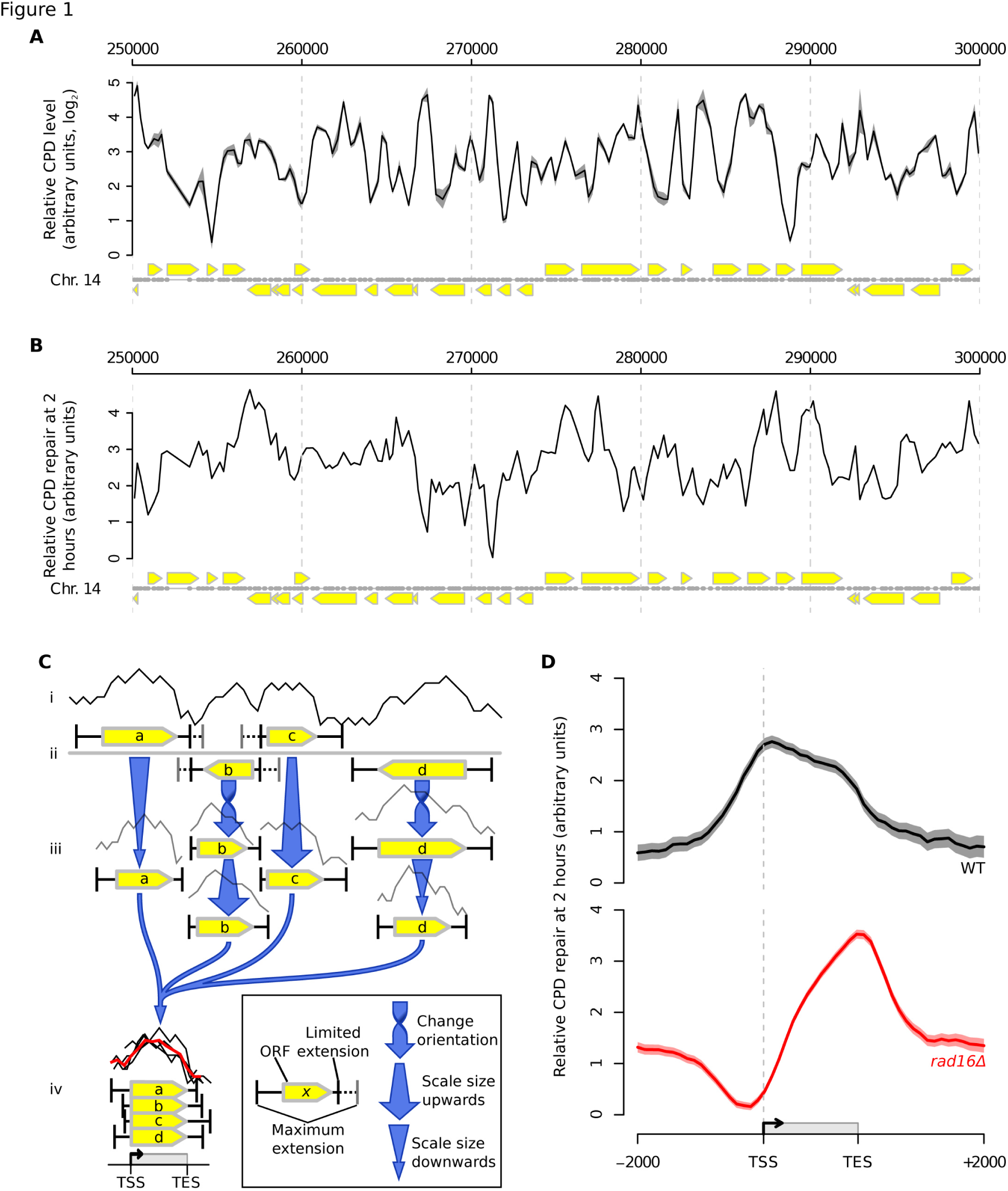
Genome-wide UV-induced DNA repair is organised around gene structure throughout the genome. (**A**) *A linear genome plot of a section of chromosome 14 showing 3D-DIP-Chip results from wild-type cells*. The black line shows the mean (n = 3) CPD level observed immediately after UV irradiation (100 J/m^2^). The shading highlights the SEM. Grey dots indicate the positions of microarray probes. Yellow arrows indicate ORF positions and their direction of transcription. CPD levels are plotted as arbitrary units on the y-axis. (**B**) *A linear genome plot of a section of chromosome 14 showing CPD repair rates*. The black line shows the mean of CPD levels 120 minutes post-UV (n = 2) subtracted from the mean at 0 minutes post-UV shown in (A). Annotations are as described in (A). (**C**) *Representation of the Sandcastle process used to overlay data from multiple ORFs to create composite plots*. A section of (i) data and (ii; labelled a, b, c, and d) ORFs are shown. Bars extending from the ORFs show the maximum length of extensions to incorporate surrounding data. Where these extensions overlap with adjacent ORF extensions (dotted line sections) they are both limited to a point halfway between the two, such that any given data point is plotted no more than once. (iii) Sections of data are then all oriented in the same direction and scaled up-or downwards so that each ORF covers the same length. (iv) These are then overlaid and an average of all the data sections calculated (red line). For all plots shown here ORFs from 500 to 1,500 bp in size were analysed (n = 2,974). (**D**) *Relative rates of CPD repair around ORF structures*. Solid lines show the mean of CPD repair rates in wild-type (n = 3, black line) and *rad16Δ* cells (n = 2, red line). Repair rates are calculated by subtracting the CPD levels detected 2 hours after UV irradiation from those levels detected immediately after. Shaded areas indicate the SEM, with CPD levels plotted as arbitrary units on the y-axis.

Presenting wild-type repair data in these composite plots reveals a uniform distribution of repair rates in the intergenic regions of the genome flanking the ORFs, with a gradual increase in repair rates in the promoter regions of genes, reaching a peak repair rate at transcription start sites (Figure 1D; black line). Enhanced rates of repair are then observed throughout the ORFs, with rates dropping to basal levels at transcription end sites (TES). It has previously been established that the enhanced rate of repair in ORFs is due to the effects of TC-NER operating in actively transcribing genes. To confirm this, we analysed CPD repair rates in the 15% of genes being expressed at the lowest levels in the genome. We recently reported global gene expression data in wild-type cells either in untreated or irradiated with UV (Zhou et al. 2015). We identified the genes expressing the lowest levels of transcription following UV damage and plotted the relative rates of CPD repair in these genes and compared these repair rates to the rates observed in the remaining 85% of the genome. Supplementary Figure S1 shows that in the 15% of genes showing the lowest level of transcription, no enhanced rates of repair in the ORF is observed compared to the higher rates of repair observed in ORFs in the remaining 85% of the genome that are transcribed at higher rates.

To examine the contribution of GG-NER to the above patterns of CPD repair rates, we examined events in *RAD16* deleted cells. In the absence of Rad16 there is a marked reduction in rates of repair around promoter regions, with a gradual increase within ORFs before reducing again in the downstream regions (Figure 1D; red line). This altered pattern is due to the absence of the GG-NER pathway, resulting in the loss of repair of the non-transcribing DNA. It is important to note that the rates of repair we present do not reflect absolute levels of lesion removal; rather they show the distribution of relative repair rates at sites throughout the genome. For this reason we plotted repair data from different mutants separately and with arbitrary units on the y-axis, as shown in Figure 1D. These results demonstrate that the GG-NER complex plays an important role in generating the pattern of DNA repair rates observed in wild-type cells, and suggest that the repair mechanism may be organised within the genome. We next considered the mechanism by which this GG-NER organisation is achieved in the genome of wild-type cells.

### GG-NER is organised and initiated from Abf1 binding sites found at thousands of locations in the yeast genome

We previously demonstrated that the yeast GG-NER apparatus is a heterotrimeric complex comprised of Abf1, Rad7 and Rad16 (Reed et al. 1999). Abf1 has a wide range of functions in processes including transcription (Buchman et al. 1988; Miyake et al. 2004; Yarragudi et al. 2007; Schlecht et al. 2008), gene silencing (Boscheron et al. 1996; Zou et al. 2006; Zhang et al. 2012), replication (Rhode et al. 1992) and NER (Yu et al. 2004; Yu et al. 2009). We have reported that binding of the Abf1 component of the GG-NER complex to one of its DNA recognition sequences promotes efficient GG-NER both *in vitro* and *in vivo* (Yu et al. 2009). Using standard chromatin immunoprecipitation (ChIP) and qPCR, we demonstrated Abf1 binding at a single Abf1 consensus binding site called the ‘I silencer’ located at the yeast *HML alpha* locus (Yu et al. 2009). Mutation of this DNA consensus site caused loss of Abf1 and GG-NER complex binding, resulting in a domain of reduced GG-NER efficiency extending from the mutated Abf1 DNA binding site. These data suggested that, prior to UV damage, the GG-NER complex might be localised at specific genomic locations, defined by the Abf1 component of the GG-NER complex binding to its consensus DNA binding motif. To determine whether other Abf1 binding sites represent locations from which domains of GG-NER are organised and initiated in response to UV damage, we used ChIP-chip and the Sandcastle software package to measure the chromatin occupancy of each component of the GG-NER complex before and after DNA damage. We first measured genome-wide Abf1 binding and performed peak detection, which found around 3,800 sites distributed throughout the yeast genome. An example of a linear genomic plot of Abf1 binding in a section of chromosome 14 is shown in Supplementary Figure S2. Other workers have investigated Abf1 binding using a variety of different methods (Yarragudi et al. 2007; Ganapathi et al. 2011; Kasinathan et al. 2014) and our study found similar binding profiles to the most recently reported study, which employed a next-generation sequencing (NGS) based method (Zentner et al. 2015). We demonstrate that these sites are located predominantly in intergenic regions, mainly in promoter regions of genes and to a lesser extent at TES (Figure 2A). Plotting Abf1 at its binding sites shows no marked change in response to UV, exhibiting only a slight reduction in overall binding 30 minutes after UV irradiation (Figure 2B). To assess the genomic distribution of Abf1 in more detail, we plotted its binding in relation to gene structure. This shows that Abf1 is highly enriched in promoter proximal regions (Figure 2C, black line). Within ORFs Abf1 occupancy is detected at much lower levels, while elevated levels of enrichment are detected downstream of the TES. These results show that Abf1 occupancy and its overall distribution in relation to ORF structure do not change markedly immediately after UV irradiation (Figure 2C, dark grey line). A small loss in overall Abf1 occupancy, which is evenly distributed across the composite gene plots, is detected at 30 minutes after UV irradiation (Figure 2C, light grey line). We conclude that Abf1 is primarily bound at promoters and other intergenic, non-transcribed regions of the genome, and Abf1 occupancy in chromatin is stable in response to UV irradiation.

**Figure 2.**
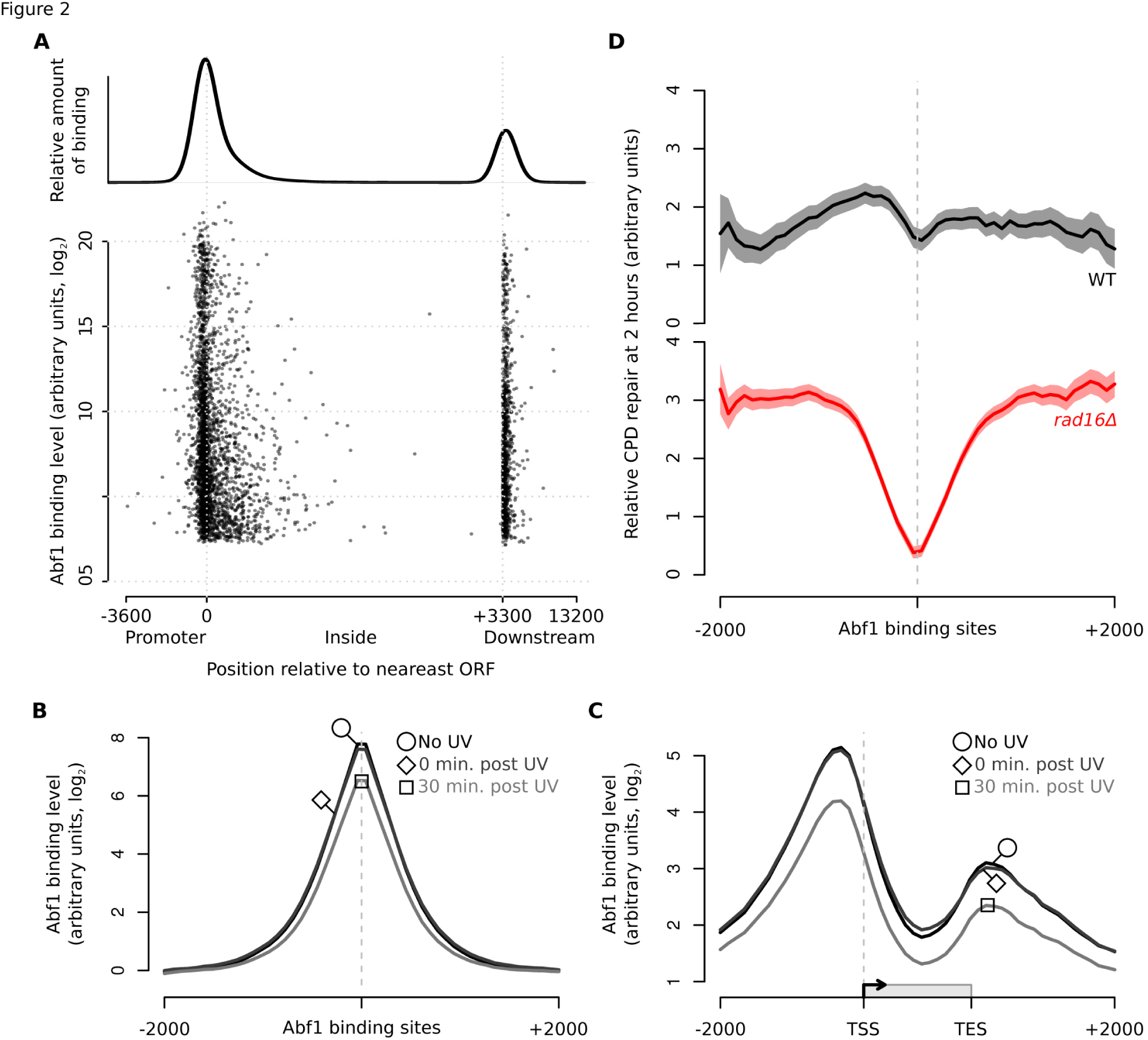
GG-NER is organised from ABF1 binding sites and Abf1 binding does not change significantly in response to UV. (**A**) *The positions of Abf1 binding relative to ORFs*. Abf1 binding levels at the ~3,800 detected binding sites are shown. Each binding site is represented by a single data point, with the overall relative amount of binding throughout the region shown above. (**B**) *ChIP-Chip data for Abf1 binding*. Data is shown for unirradiated (black, circle highlight), 0 min post-UV (dark grey, diamond highlight), and 30 min post-UV (light grey, square highlight) datasets. Solid lines show the means of 3 datasets per time point. (**C**) As (B) plotted around ORF structure (see Figure 1C). (**D**) *Relative CPD repair rates around Abf1 binding sites*. The data depicted in Figure 1D is used here to plot the relative rates of CPD removal around Abf1 binding sites in wild-type (black) and *rad16Δ* cells (red). Solid lines show the mean of CPD repair rates in wild-type (n = 3, black line) and *rad16Δ* cells (n = 2, red line). Repair rates are calculated by subtracting the CPD levels detected 2 hours after UV irradiation from those levels detected immediately after. The shaded areas show the SEM, while CPD levels are plotted as arbitrary units on the y-axis.

To determine whether GG-NER is organised from these Abf1 binding sites we plotted our DNA repair rate data for both wild-type and GG-NER defective *RAD16* deleted cells as composite plots centred on the detected Abf1 binding sites (Figure 2D upper and lower panels respectively; black and red lines). This revealed that the rates of repair in *rad16* deleted cells are markedly reduced around Abf1 binding sites compared to *RAD16* wild-type controls. Importantly, plotting the distribution of DNA repair rates at an equal number of randomly generated simulated ORFs, reveals an even distribution of repair rates in both wild-type and *rad16* deleted cells (Supplementary Figure S3). These observations confirm that Abf1 binding sites play a significant role in orchestrating GG-NER in the genome.

### The GG-NER complex protein Rad7 localises to Abf1 binding sites

Our previous studies examining events at the *HML alpha* locus (Yu et al. 2009) indicated that the GG-NER complex binds to chromatin at this Abf1 binding site, where it promotes efficient DNA repair. We considered the possibility that GG-NER is organised in the genome by the GG-NER complex localising at multiple Abf1 binding sites, priming the complex for efficient GG-NER throughout the genome. To investigate this, we used ChIP-chip to measure the genome-wide occupancy of Rad7 and plot the data at Abf1 binding sites, which reveals a strong enrichment of Rad7 occupancy at these sites (Figure 3A; black line). This extends our previous observations (Yu et al. 2009), demonstrating that Rad7 co-localises at multiple Abf1 binding sites throughout the genome in the absence of UV damage. Next, we investigated the effect of UV irradiation on Rad7 binding in wild-type cells. Following UV irradiation, after 15 minutes there is a marked reduction of Rad7 localisation at Abf1 binding sites, but not complete loss of its occupancy from chromatin (Figure 3A; grey line). Displaying the data as composite gene plots orientates the Abf1 binding sites in relation to ORFs (Figure 3B). This enables the UV-induced redistribution of Rad7 to be discerned over the whole ORF structure. This reveals that in response to UV irradiation Rad7 dissociates from Abf1 binding sites in promoter and downstream regions and redistributes predominantly into ORFs and upstream promoter regions.

**Figure 3.**
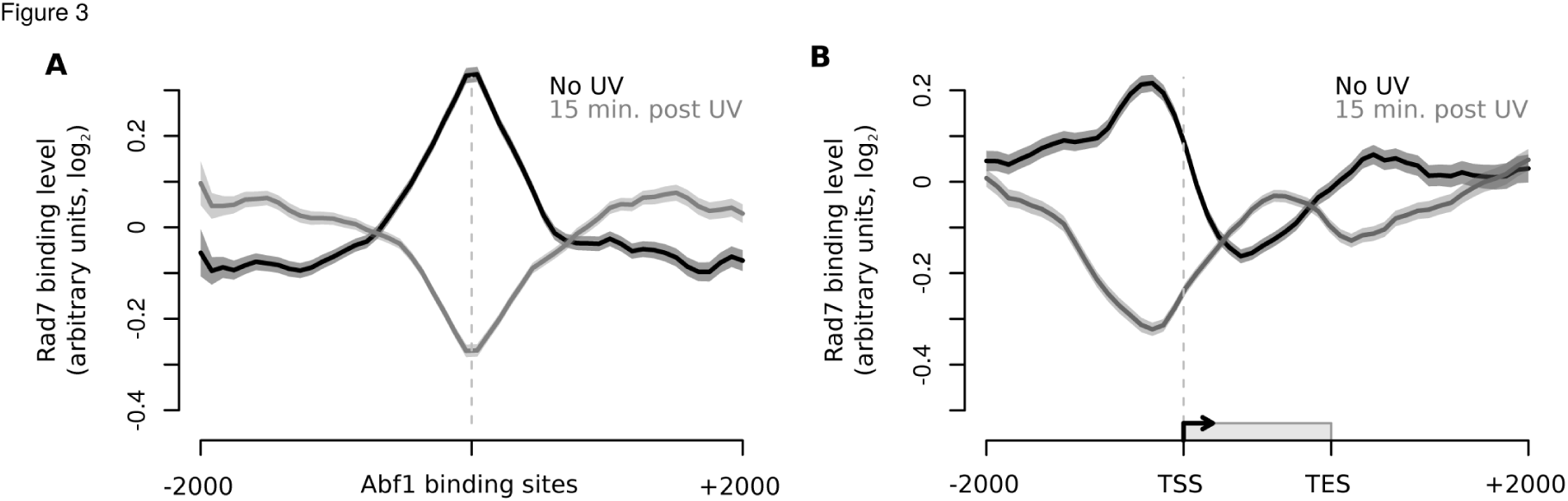
The co-localisation of the GG-NER factor Rad7 in chromatin at Abf1 binding sites and its redistribution in response to UV irradiation is similar to Rad16. (**A**) Rad7 binding data around ORF structure for unirradiated (black) and 30 min post UV (grey) datasets. Solid lines show the means of three datasets per time point and shaded areas show the SEM. (**B**) As (A) plotted around detected Abf1 binding sites.

### The Rad7 and Rad16 proteins co-localise with Abf1 in the genome

Rad7 and Rad16 form two of the core components of the GG-NER complex and their protein-protein and genetic interactions, as well as their phenotypes are well established (Verhage et al. 1994; Reed et al. 1998). To determine whether the genomic occupancy of Rad16 is similar to that of Rad7 and Abf1, we performed ChIP-Chip for Rad16 and plotted the resulting data around Abf1 binding sites (Figure 4A, black and grey lines). We note the loss of occupancy from Abf1 binding sites as observed for Rad7 (Figure 3A). Similar observations are made when examining events as composite gene plots (Figure 4B). Slightly reduced levels of Rad16 chromatin occupancy are observed at the 30 minute time point after UV irradiation. As anticipated, Rad16 distribution around ORFs prior to UV irradiation is very similar to that of Abf1 (Figure 2C, black line) and Rad7 protein binding (Figure 3B, black line), showing enrichment in intergenic regions, especially at promoters upstream of the TSS, and lower levels of enrichment downstream of TES (Figure 4B, black line). Notably, the redistribution we observe for Rad16 30 minutes after exposure to UV damage is very similar to that observed for Rad7, namely a loss of occupancy from their initial binding positions and increased accumulation in upstream regions of the promoter and within the ORFs (compare Rad7, Figure 3B, grey line, with Figure 4B, grey line). Finally, we established that the distribution of Rad7 is dependent on the GG-NER complex by performing ChIP-Chip for Rad7 in a *RAD16* deleted strain. As shown in Supplementary Figure S5A, displaying the data as composite gene plots reveals that the normal pattern of Rad7 distribution prior to UV irradiation depends on *RAD16*. In this mutant background Rad7 is distributed throughout the ORFs prior to UV irradiation and is not redistributed 15 minutes after UV irradiation (Supplementary Figure 5A, compare light and dark red lines). These observations were confirmed when we examined events in the context of GG-NER complex occupancy at Abf1 binding sites (Supplementary Figure S5B). Collectively, these results demonstrate that prior to UV irradiation the Rad7 and Rad16 components of the GG-NER complex are located predominantly at Abf1 binding sites found in intergenic regions of the genome, particularly in promoter and downstream regions of genes. In response to UV damage, a complex of Rad7 and Rad16 is redistributed away from Abf1 binding sites and occupies locations within the ORFs.

**Figure 4.**
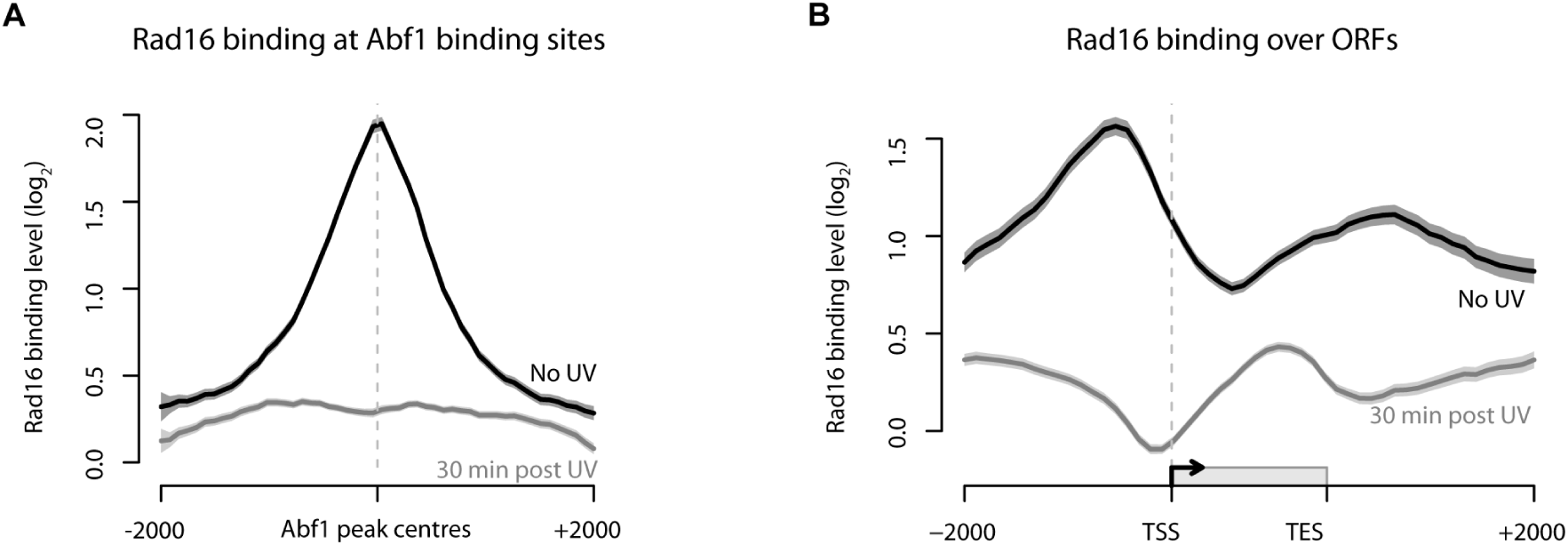
Rad16 associates with chromatin surrounding Abf1 binding sites and its redistribution in response to UV. (**A**) Rad16 binding data around each of the detected Abf1 binding sites for unirradiated (black) and 30 min post UV (grey) datasets. Solid lines show the means of three datasets per time point and shaded areas show the SEM. (**B**) As (A) plotted around ORF structure.

### Rad16 genomic occupancy depends on its ATPase and RING domain functions

We next investigated which Rad16 activities are responsible for the distribution of Rad16 observed both before and after UV radiation. Rad16 contains within its structure two functional regions that contribute to efficient GG-NER: two SWI/SNF ATPase domains; and an E3 ubiquitin ligase RING domain (Figure 5A). It has previously been reported that mutationally inactivating these domains individually reduces repair rates and results in UV sensitivity intermediate between those of wild-type and *RAD16* deletion strains, while mutating both domains together generates UV sensitivity equivalent to that of *rad16* null strains (Ramsey et al. 2004; Yu et al. 2011). To determine the contributions of these domains to the initial distribution and UV-induced redistribution of Rad16 we used ChIP-Chip to measure its occupancy in strains containing point mutations in the ATPase domain, the RING domain, or both domains together. Strains expressing these mutated genes produce full-length Rad16 proteins (Supplementary Figure S4A) that can associate with chromatin (Supplementary Figure S4B), as shown by western blot analysis. The distributions of Rad16 at Abf1 binding sites before, and after UV-irradiation in wild-type cells (Figure 4A) are lost in the ATPase/RING double mutant strain (Figure 5B; dark and green solid lines respectively). Similar results are seen in the composite gene plot (Figure 5C), confirming the loss of the wild-type pattern of Rad16 distribution. These data show that the distribution of Rad16 before, and its redistribution after UV irradiation depends on functional ATPase and RING domains.

**Figure 5.**
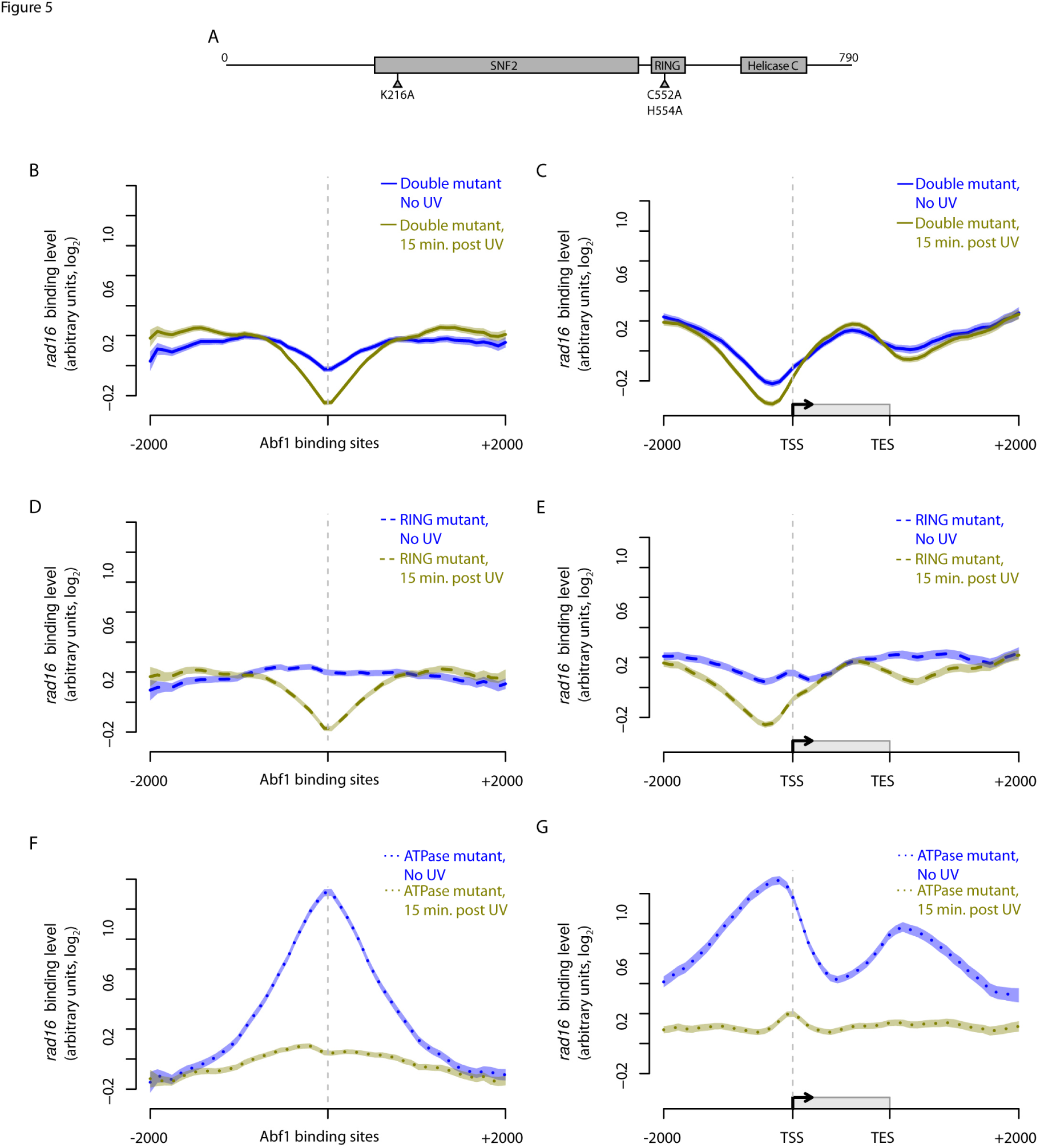
The activity of both the ATPase and RING domain of Rad16 determine its chromatin occupancy before and after UV irradiation. (**A**) *Representation of the linear structure of Rad16*. The amino acids targeted by the point mutations introduced in the ATPase (K216A) and RING domains (C552A, H554A) are highlighted. (**B** – **G**) *Composite plots of Rad16 chromatin occupancy in the mutants described*. Mutated Rad16 binding data around ABF1 binding sites and ORF structures in the absence of UV irradiation (dark blue) and 15 minutes after UV irradiation (green). The Rad16ATPase/RING double mutant binding data (solid lines) are shown in B (around ABF1 binding sites) and C (around ORFs). The binding data for Rad16 RING domain mutant (dashed lines) is shown in D (around Abf1 binding sites) and E (around ORFs). Finally, the Rad16ATPase mutant data (dotted lines) is depicted in F (around Abf1 binding sites) and in G (around ORFs). Lines show the means of three datasets per condition and shaded areas show the SEM.

Notably, we found that inactivating Rad16 E3 ligase function by mutating its RING domain, results in the loss of its wild-type occupancy at Abf1 binding sites in the absence of UV irradiation, taking up an even distribution in unirradiated cells (Figure 5D and E, dark blue dashed line). However, in response to UV, some redistribution of this mutant Rad16 protein still occurs, resulting in a small change in its distribution (Figure 5D and E; compare dark blue lines with green lines). This indicates that when the ATPase domains are intact, some UV-induced change in the distribution of Rad16 can still occur. Examining the same events in the ATPase mutant Rad16 strain shows a distribution in the absence of UV damage similar to that of the wild-type strain (Compare Figure 4A and B, with Figure 5F and G; dark blue dotted lines). A large reduction in the occupancy of Rad16 from its initial position is observed in response to UV irradiation, but not showing the redistributed pattern into the ORFs observed in wild-type cells (Figure 5F and G, green lines). We conclude from this that Rad16 E3 ubiquitin ligase activity associated with the RING domain of Rad16 is required for establishing and maintaining Rad16 binding at Abf1 binding sites prior to UV irradiation, while the ATPase activity of Rad16 is dispensable. However, the ATPase domain is required for Rad16 redistribution after UV radiation. Therefore, both Rad16 activities function in concert to contribute to the normal occupancy of Rad16 in chromatin before, and its redistribution after, UV irradiation.

### The GG-NER complex regulates genome-wide distribution of Gcn5 chromatin occupancy before and after UV irradiation

UV-induced chromatin modifications contribute to efficient repair at the *MFA2* locus during GG-NER, through Gcn5-dependent hyperacetylation of histone H3K9 and H3K14 (Yu et al. 2005). In our previous work we noted that UV-induced acetylation occurs independently of the core NER factors Rad4 and Rad14, demonstrating that functional NER is not required for this activity. However, we found that the *RAD7* and *RAD16* genes are required for UV-induced acetylation of histone H3K9/K14 at the *MFA2* locus and that this was achieved by the GG-NER complex controlling chromatin occupancy of the HAT Gcn5 (Yu et al. 2005). We also reported that this process promotes chromatin remodelling, making the chromatin more accessible to restriction enzyme digestion (Yu et al. 2011). To determine how the GG-NER complex controls Gcn5 occupancy within the genome, we performed ChIP-Chip experiments for Gcn5 and analysed the genome-wide chromatin binding data. This established that Gcn5 is enriched at Abf1 binding sites prior to UV irradiation (Figure 6A, black line, circle highlight), similar to the components of the GG-NER complex (Figures 3A and 4A). Gcn5 occupancy in the vicinity of these sites increases immediately following UV irradiation (Figure 6A, dark grey line, diamond highlight), and gradually reduces after 15 minutes (Figure 6A, mid-grey line, square highlight), with further reduction in Gcn5 occupancy observed 60 minutes after UV irradiation (Figure 6A light-grey line, triangle highlight). Importantly, enrichment of Gcn5 in the vicinity of Abf1 binding sites is retained during this period (Figure 6A). To investigate whether the GG-NER complex plays a role in regulating the UV-induced change in Gcn5 occupancy, we measured genome-wide Gcn5 binding in the absence of Rad16. The results show that Gcn5 binding at Abf1 binding sites prior to UV irradiation is similar to that seen in wild-type cells, but at slightly lower levels (Figure 6B, solid red line, circle highlight). We observed an initial rapid UV-induced recruitment of Gcn5 to Abf1 binding sites in the absence of Rad16, but at levels lower than observed in wild-type cells, indicating the importance of the GG-NER complex for wild-type levels of Gcn5 recruitment (Figure 6B, dark pink line, diamond highlight). However, 15 minutes after UV irradiation Gcn5 is no longer enriched at these sites in the *RAD16* deleted strain compared to wild-type cells (Figure 6B, mid pink line, square highlight), and occupancy is even further reduced after 60 minutes (Figure 6B, light pink line, triangle highlight). This result clearly shows that in addition to regulating the initial UV-induced level of Gcn5 occupancy, Rad16 is required for maintaining Gcn5 occupancy on chromatin in the vicinity of Abf1 binding sites at later time points following UV irradiation. Figure 6C reveals a similar Gcn5 distribution to that of Abf1 and the GG-NER factors prior to UV irradiation. This pattern manifests as a strong promoter proximal enrichment and lower levels of enrichment at the 3’ ends of genes. Figure 6D shows that in *RAD16* deleted cells Gcn5 occupancy is reduced, predominantly in the vicinity of the promoter proximal Abf1 binding sites compared to wild-type cells at the 15 minute and 1 hour time points. These observations demonstrate the role of the GG-NER complex in regulating the retention of the HAT Gcn5 on the chromatin in response to UV radiation. To further explore the role of the GG-NER complex in retaining Gcn5 occupancy on the promoter proximal chromatin regions in response to UV radiation, we plotted the data for Gcn5 binding in wild-type and *RAD16* deleted cells at each of the different time points measured (Supplementary Figure S6A-D). We conclude from these results that the GG-NER complex regulates the wild-type recruitment and retention of Gcn5 occupancy in chromatin in promoter proximal domains prior to and following UV irradiation.

**Figure 6.**
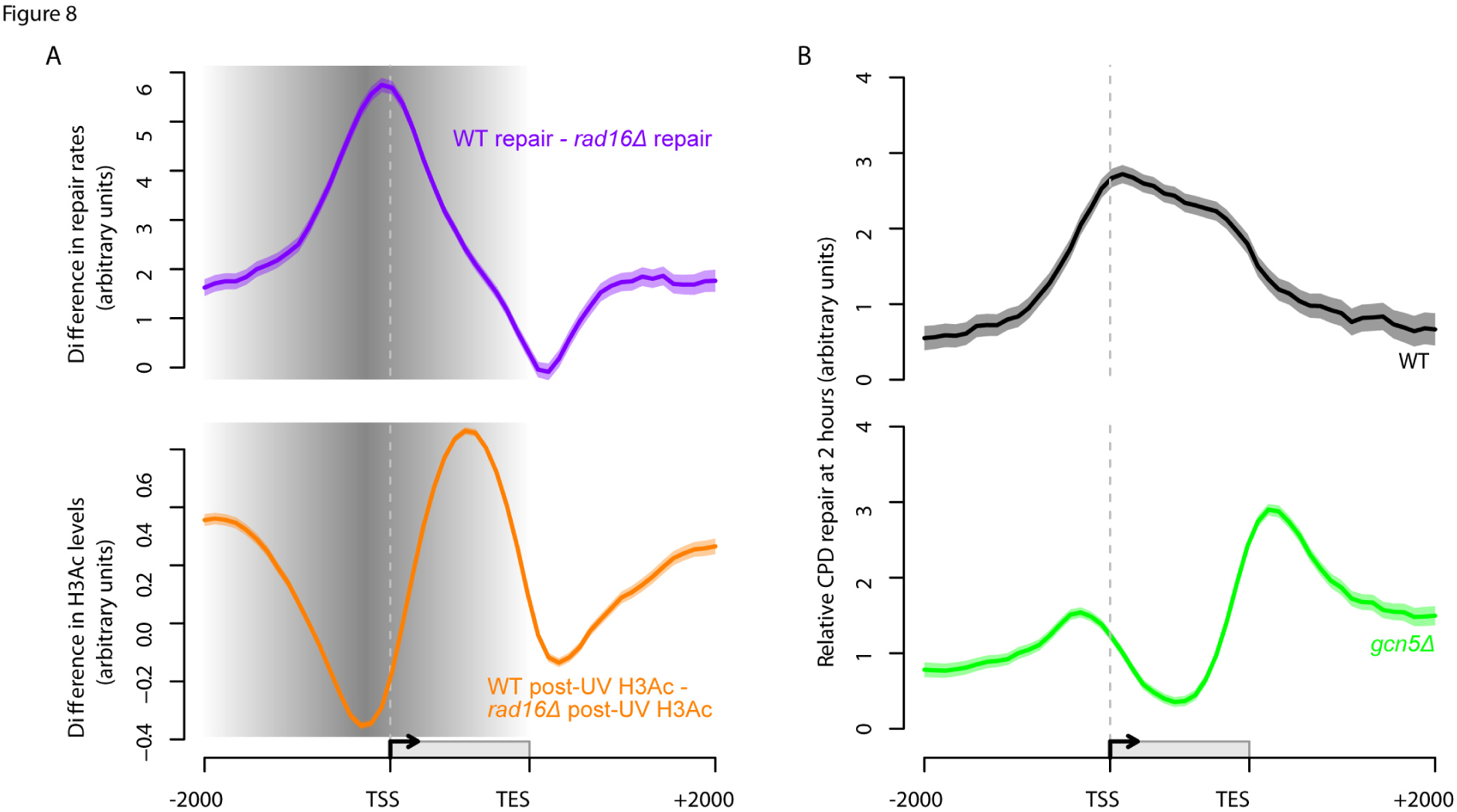
The GG-NER pathway coordinates lesion removal by controlling UV-induced histone H3 acetylation in genomic domains around ABF1 binding sites. (**A**) Rad16-dependent repair (purple line) is calculated by subtracting *rad16Δ* repair from the wild-type repair data. UV-induced H3Ac (orange line) is calculated by subtracting the *rad16Δ* post-UV H3Ac from the wild-type data. The shading highlights the domain where lesion removal and UV-induced H3 acetylation are controlled by the GG-NER complex, initiated predominantly from sites of Abf1 binding. (**B**) Relative rates of CPD removal around ORF structures in wild-type (n = 3, black) and *gcn5Δ* (n = 2, green) cells. Solid lines show the mean of relative CPD repair rates levels, with the shaded areas highlighting the SEM. CPD levels are plotted as arbitrary units on the y-axis.

### The GG-NER complex regulates the UV-induced genomic distribution of histone H3 acetylation

We previously demonstrated that the UV-induced chromatin remodelling necessary for GG-NER is promoted by the induction of histone H3 acetylation at the *MFA2* gene. We established that this is controlled by the GG-NER complex, which regulates occupancy of the HAT Gcn5 on the chromatin at this locus (Yu et al. 2011). Having established the genomic distribution of Gcn5 occupancy in chromatin before UV irradiation and the change in its distribution afterwards, we next investigated how this epigenetic histone modification catalysed by this HAT is distributed within the genome and how it changes in response to UV irradiation. In wild-type cells we observe a distinctive ‘m-shaped’ pattern for this epigenetic marker around Abf1 binding sites (Figure 7A, black line). Histone H3 acetylation reaches a maximum at sites approximately 300 bp either side of Abf1 binding sites and reduces at positions located further away. The lower levels of histone H3Ac centred at Abf1 binding sites is likely caused by the absence of histones at these predominantly nucleosome free regions (NFRs) (Hartley and Madhani 2009; Ozonov and van Nimwegen 2013). In response to UV irradiation an increase in histone H3 acetylation is detected, with a maximum enrichment observed at approximately 500 bp on either side of the Abf1 binding sites (Figure 7A, grey line) and the characteristic ‘m-shaped’ pattern of histone modification is retained. However, in a *RAD16* deleted strain, lower levels of histone H3 acetylation are observed in the absence of UV irradiation compared to wild-type cells (Figure 7A, dark red line), in line with the reduced Gcn5 occupancy we observed previously (Figures 6A and C). This indicates that Rad16 plays a role in determining the basal level and distribution of histone H3 acetylation in the absence of DNA damage. In response to UV irradiation, induction of histone H3Ac can still be observed (Figure 7A, light red line), corresponding to the Rad16-independent recruitment of Gcn5 to Abf1 binding sites described in the previous section (Figures 6A and B). We note that Rad16-dependent UV-induced increase in histone H3 acetylation observed in wild-type cells corresponds to the redistribution of the GG-NER complex components Rad7 (Figure 3B) and Rad16 (Figure 4B) away from their initial positions at Abf1 binding sites. Importantly, the UV-induced distribution of histone H3Ac around Abf1 binding sites and ORFs (Figure 7B, shaded areas) is significantly different in *RAD16* deleted cells compared to wild-type cells. By contrast, in the absence of UV irradiation, H3Ac is distributed around genes in a similar fashion to the occupancy of Gcn5 in wild-type cells (compare Figure 7B, black line with Figure 6C, black line). Similarly, the distribution of histone H3 acetylation and Gcn5 occupancy observed in *rad16Δ* cells is comparable, albeit at lower levels than in wild-type cells (compare Figure 7B, red line with Figure 6D, red line). In response to UV irradiation histone H3 acetylation is induced in both wild-type and *rad16Δ* cells (Figure 7B, grey and light red lines respectively). However, the UV-induced pattern of histone H3 acetylation is different in *rad16Δ* cells compared to the wild-type. The shaded area in Figure 7B identifies the GG-NER complex dependent histone H3 acetylation in response to UV irradiation. The altered pattern of histone H3Ac in the *RAD16* deleted strain extends from within the promoter regions at Abf1 binding sites throughout the ORFs and also into upstream intergenic regions (Figure 7B, grey line). We conclude that the GG-NER complex directs the UV-induced propagation of histone H3 acetylation by regulating the residency of Gcn5 in the genomic domains described.

**Figure 7.**
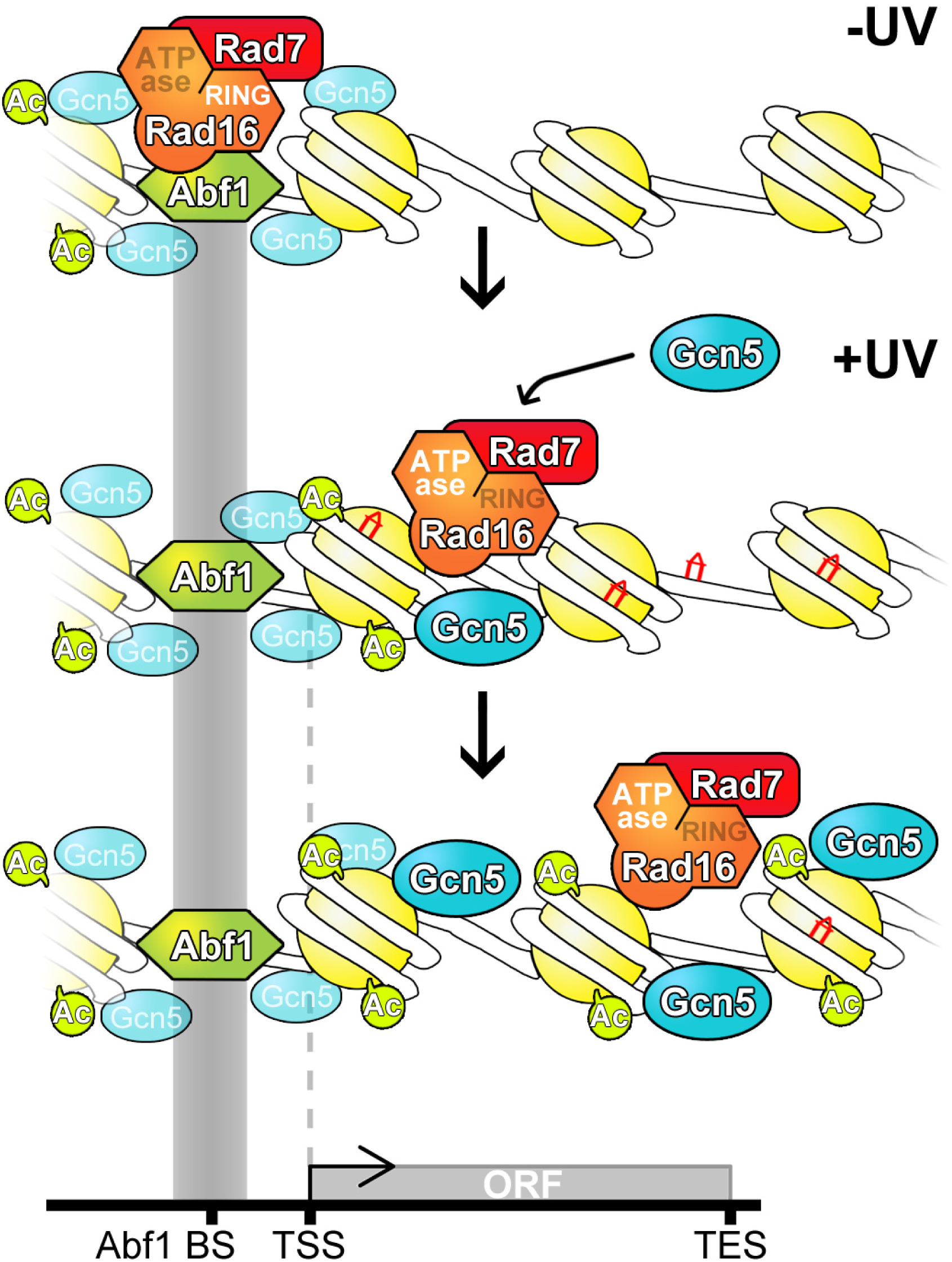
Model to illustrate how GG-NER is organised in the yeast genome. Top panel: In undamaged cells the GG-NER complex is located at multiple ABF1 binding sites predominantly in the promoter regions of genes. This occupancy is dependent on the RING domain of the Rad16 protein. The enrichment of GG-NER-independent basal levels of Gcn5 can be detected at these sites. Middle panel: In response to UV irradiation, the GG-NER complex dissociates from the Abf1 component at ABF1 binding sites. This process depends on the activity of the ATPase domain in Rad16. Concomitantly, the HAT Gcn5 is recruited onto the chromatin with its increased levels and distribution dependent on the Rad7-Rad16 GG-NER complex. Bottom panel: During this process histone H3 acetylation is increased over a domain defined by the redistribution of the Rad7-Rad16 proteins from ABF1 binding sites. This mechanism drives the chromatin remodelling necessary for the efficient repair of UV damage.

### Defective UV-induced chromatin remodelling results in domains of altered DNA repair rates distributed throughout the genome

We previously reported that UV-induced histone H3 acetylation promotes chromatin remodelling necessary for efficient GG-NER (Yu et al. 2005). In the present study we have shown how the GG-NER complex regulates this process, and how these events are organised within the yeast genome. In Figure 1D we demonstrated the effect on the distribution of genomic DNA repair rates when the GG-NER pathway is abrogated in *rad16* mutated cells. In Figure 8A (upper panel, purple line) we have plotted the difference between the wild-type and Rad16 mutant repair rates to define the genomic regions most affected by loss of the GG-NER pathway. This shows that the biggest effect on DNA repair rates in the absence of GG-NER occurs in promoter regions, just upstream of transcription start sites, where Abf1 binding sites are predominantly located, and extend in both directions into the upstream promoter as well as into the ORF regions. A similar analysis plotting the difference between UV-induced histone H3 acetylation in wild-type and *RAD16* deleted cells (Figure 8A, lower panel, orange line) reveals the reciprocity between repair rates and UV-induced histone H3 acetylation levels in the absence of GG-NER, and defines the genomic domains from which GG-NER organises and initiates chromatin remodelling and repair, highlighted by grey shading. Strikingly, the genomic regions that exhibit defective histone H3 acetylation in GG-NER-defective cells align with the regions of altered DNA repair rates observed in these cells (Figure 8A). To probe the central importance of histone H3 acetylation on the distribution of genomic DNA repair rates, we deleted *GCN5* and showed that this results in complete loss of UV-induced histone H3Ac at K9/K14 observed in wild-type cells (Supplementary Figure S7). This demonstrates the central role of this chromatin modifier in promoting the wild-type distribution of UV-induced histone H3Ac within the genome. Finally, we measured removal of UV-induced DNA damage in the absence of Gcn5. Figure 8B compares the relative repair rates in *GCN5* deleted cells (lower panel, green line) to those in wild-type cells (upper panel, black line) in the context of gene structure. Importantly, these data establish that the distribution of DNA repair rates is completely disrupted in the absence of the HAT Gcn5. This confirms the importance of the GG-NER complex in regulating the UV-induced, Gcn5 catalysed histone H3 acetylation on the wild-type distribution of genomic DNA repair rates.

**Figure 4.**
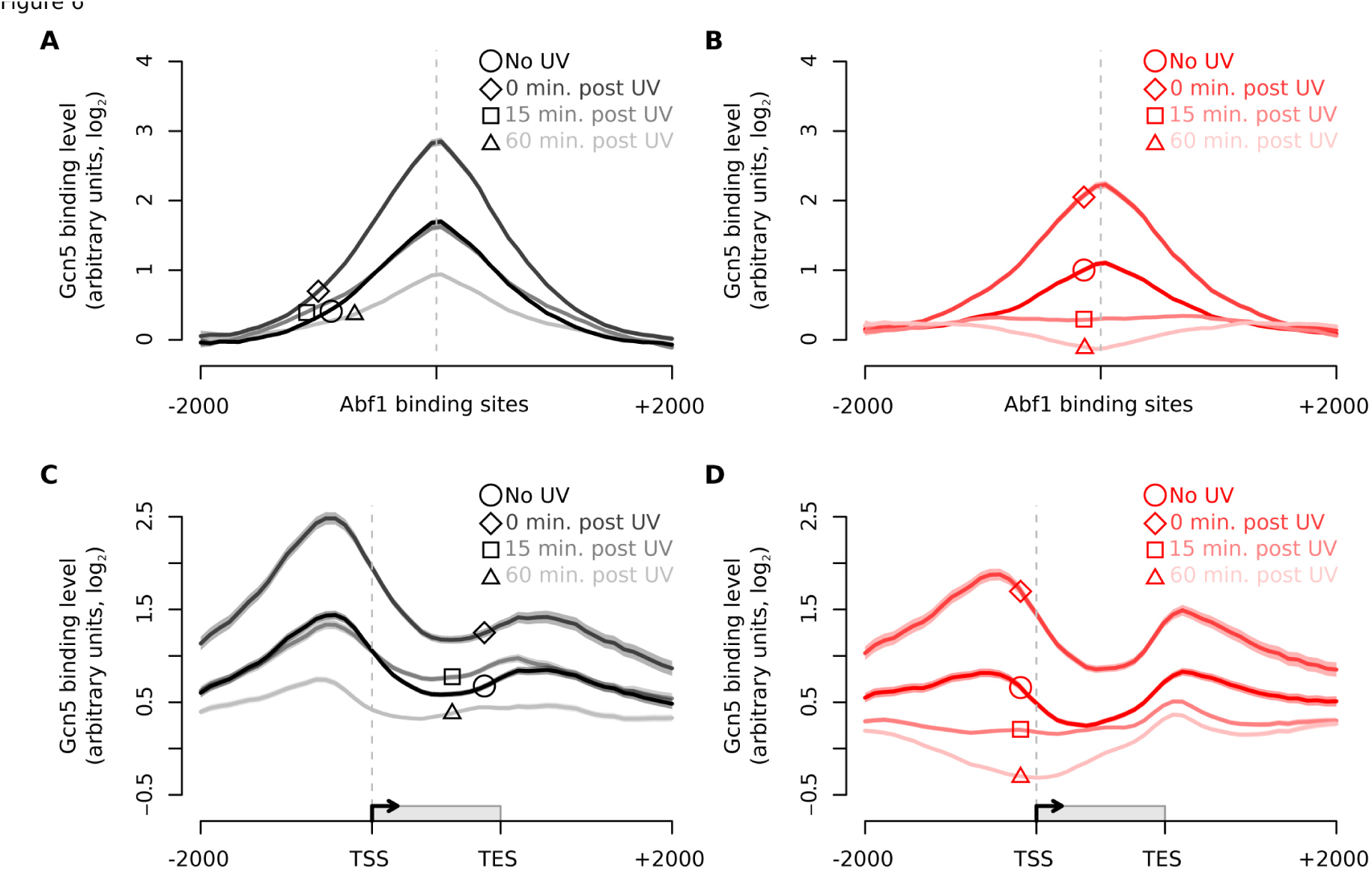
Gcn5 is recruited to ABF1 binding sites and ORFs in response to UV in a Rad16-dependent manner. (**A**) Gcn5 binding data in wild-type cells around each of the detected ABF1 binding sites for unirradiated (black; circle highlight), 0 min post-UV (dark grey; diamond highlight), 15 min post-UV (mid grey; square highlight) and 30 min post UV (light grey; triangle highlight) datasets. Solid lines show means (n = 3, 3, 2 and 3 respectively) and shaded areas show the SEM. (**B**) Gcn5 binding data in *rad16Δ* cells around each of the detected ABF1 binding sites for unirradiated (red; circle highlight), 0 min post-UV (dark pink; diamond highlight), 15 min post-UV (mid pink; square highlight) and 30 min post UV (light pink; triangle highlight) datasets. Solid lines show the means of two datasets per time point and shaded areas show the SEM. (**C**) As (A) plotted around ORF structure (see Figure 1C). (**D**) As (B) plotted around ORF structure (see Figure 1C).

## Discussion

In this report, we have provided new insights into understanding the processes that govern genome stability and how these events are organised within the genome. In previous studies we showed (Reed et al. 1999; Yu et al. 2009) that a single Abf1 binding site located at the *HML* alpha locus functions as a site from which DNA repair is initiated (Yu et al. 2009). Therefore, we considered the possibility that the structure of the genome is organised in such a way that ensures the efficient rate of removal of DNA damage by the GG-NER pathway. We speculated that Abf1 binding sites in general might represent regions from which GG-NER is organised to promote efficient DNA repair of chromatin in the genome. To investigate this possibility, we embarked herein to map genomic rates of DNA repair in relation to the genomic occupancy the GG-NER complex components, both before and after exposing cells to UV irradiation. We focused on repair of UV-induced cyclobutane pyrimidine dimers by the GG-NER pathway in yeast. Using 3D-DIP-Chip (Teng et al. 2011; Powell et al. 2015), we generated genome-wide DNA damage and repair profiles both in wild-type and mutant yeast strains. This showed that in wild-type cells, both the initial pattern of CPD induction and the subsequent pattern of their relative DNA repair rates, appear to be heterogeneously distributed throughout the genome when viewed as a linear representation of the chromosomes. Importantly however, presenting such data as composite gene plots around ORFs (upper panel of figure 1D), revealed a level of organisation of genomic repair rates in wild-type cells that was previously unknown. We noted enhanced rates of CPD removal within the ORFs in wild-type cells, which is consistent with the known contribution of the transcription coupled repair pathway to the rapid removal of lesions from the transcribed strand of active genes (Mellon et al. 1987). A similar observation was made in a recent study measuring genomic DNA repair using the NGS-based method XR-seq in human cells (Hu et al. 2015; Adar et al. 2016). To examine the effect of removing the GG-NER pathway, we measured DNA repair rates in *RAD16* deleted cells, and observed a significantly altered distribution of genomic DNA repair rates (Figure 1D, lower panel). The altered genomic DNA repair rate profile represents the contribution of the TC-NER pathway, which remains intact in these mutant cells. It’s important to note, however, that the data derived from these DNA damage and repair experiments measure only the relative rates of repair and not the absolute levels of lesion removal.

We considered how the organisation of genomic DNA repair rates might be established in the genome. Thus, we examined the genomic distribution of GG-NER complex components both before and after UV treatment. Firstly, we examined Abf1 binding in the absence of UV damage and observed approximately 3,800 peaks distributed throughout the genome. The majority of these sites are located in the promoter region of genes, close to the TSS, although a second, less abundant group can be found at the 3’ end of ORFs near the TES. This demonstrates that the vast majority of Abf1 binding sites are located in intergenic, non-transcribed regions of the genome. To determine whether these sites represent locations from which GG-NER is organised, we plotted genomic DNA repair rates for wild-type cells against GG-NER defective *RAD16* deleted cells in relation to all Abf1 binding sites, which revealed significantly reduced repair in the vicinity of Abf1 binding sites in GG-NER defective cells. This demonstrates the importance of GG-NER for efficient removal of CPD lesions at these genomic locations, suggesting that GG-NER may be organised from these sites. Importantly, genomic Abf1 distribution does not change markedly in response to UV irradiation, as only a small loss of occupancy, evenly distributed across ORFs is observed. We conducted similar experiments for the Rad7 and Rad16 components of the GG-NER complex. These GG-NER components co-localise with Abf1 at multiple Abf1 binding sites in the absence of UV irradiation. This demonstrates that the GG-NER complex is chromatin bound at Abf1 binding sites in the genome in the absence of DNA damage, poising the genome for GG-NER. However, following UV irradiation and in contrast to Abf1 itself, a striking loss of Rad7 and Rad16 occupancy is seen from Abf1 sites and a distinctive redistribution of these proteins is observed extending into the ORFs away from the Abf1 binding sites. These observations demonstrate that the Abf1 component of the GG-NER complex anchors the repair factors Rad7 and Rad16 at Abf1 binding sites in the absence of DNA damage. This establishes the formation of nucleation sites for GG-NER at these genomic locations, which primes the pathway for efficient lesion removal. In response to UV irradiation, the Rad7 and Rad16 components of the complex are redistributed away from these sites, while the Abf1 component predominantly remains bound. This organisation of the genome into domains promotes the efficient removal of UV-induced DNA damage from the genome by GG-NER. Future studies will focus on the mechanism of the UV-induced dissolution of the complex.

By studying the effects of inactivating mutations in key domains of Rad16, we found that while the RING E3 ligase motif was important for the normal pre-UV irradiation distribution of Rad16 observed in wild-type cells, the ATPase domain is dispensable for this. This suggests that ubiquitylation of an as yet undefined target protein by the E3 ligase function of the GG-NER complex, is necessary for normal positioning of the complex in the genome in the absence of DNA damage. Potential candidates for such ubiquitylation include histones, which may serve to tether the GG-NER complex to the chromatin in the vicinity of Abf1 binding sites. In this regard we note that the UV-DDB complex, which is involved in GG-NER in human cells, is a component of an E3 ubiquitin ligase that ubiquitylates histone H2A in response to UV damage (Kapetanaki et al. 2006; Lan et al. 2012). In contrast, we found that the ATPase domain is required for the post-UV redistribution of Rad16 into the ORFs seen in wild-type cells. This observation is consistent with the presence of ATPase motifs in Rad16 that are required for the DNA translocase activity of the complex (Yu et al. 2004).

Our previous studies suggested that the GG-NER complex controls UV-induced histone H3 acetylation by regulating recruitment of the Gcn5 onto the chromatin (Yu et al. 2011). Examining Gcn5 occupancy on a genomic scale in our current study revealed how the GG-NER complex controls its occupancy on the chromatin at the correct genomic locations necessary to promote efficient GG-NER. In wild-type cells we observed Gcn5 occupancy in chromatin with a similar distribution to the GG-NER complex components. We noted a very rapid increase in Gcn5 occupancy in response to UV irradiation. Examining Gcn5 binding at later time points revealed a gradual overall reduction in its occupancy, but Gcn5 retention is seen in regions of the genome where we observe UV-induced GG-NER complex redistribution. We found that retention of Gcn5 at the genomic locations observed depends on UV-induced redistribution of the GG-NER complex away from its initial positions at Abf1 binding sites during a one hour period after UV damage. Consistent with a potential UV-induced direct or indirect interaction between the GG-NER complex and Gcn5, we found that the GG-NER complex controlled induction and retention of Gcn5 in response to UV at the correct genomic locations in chromatin is necessary for the normal pattern of UV-induced histone H3 acetylation (shaded areas highlighted in the context of Abf1 binding sites and composite gene plots shown in Figure 7A and B). Deletion of *RAD16* results in lower levels of histone H3 acetylation in the absence of UV damage, highlighting a role for the GG-NER complex in setting basal levels of histone H3 acetylation in the genome. Whether this affects cellular processes outside of NER, such as gene transcription, remains unknown at present. We noted that histone H3 acetylation is induced following UV irradiation in the absence of Rad16 but that in the absence of Rad16, this acetylation is not induced to wild-type levels and fails to spread towards the genomic regions where redistribution of the GG-NER complex occurs in response to UV irradiation. Finally, we compared the effect of removing the GG-NER pathway on the distribution of histone H3 acetylation, and rates of DNA repair. By plotting the difference between these wild-type and *rad16Δ* cells, we established that the regions of DNA repair most affected by loss of GG-NER correspond to the regions most affected by the UV-induced, GG-NER dependent histone H3 acetylation (Figure 8A, shaded region). We conclude that the GG-NER complex regulates the chromatin structure in the vicinity of Abf1 binding sites in response to UV irradiation by controlling the occupancy of the HAT Gcn5 on the chromatin and the UV-induced histone H3 acetylation status.

Our data lead us to a model (Figure 9) in which the Rad7 and Rad16 components of the GG-NER complex contribute to the normal distribution and steady-state levels of histone H3Ac observed in the absence of DNA damage. Following UV irradiation, Rad7 and Rad16 promote the recruitment and retention of Gcn5 to intergenic and non-transcribed regions of chromatin located within ORFs. This increased Gcn5 occupancy in turn promotes enhanced levels of histone H3 acetylation within these regions of the genome to promote the normal distribution of DNA repair rates in these genomic regions. Confirming the significance of Gcn5 mediated UV-induced histone H3 acetylation on DNA repair, we have shown that in an otherwise wild-type cell, deleting the HAT *GCN5*, which is required for the UV-induced histone H3 acetylation, completely altered the pattern of repair rates seen in wild-type cells (Figure 8B). This demonstrates the importance of the GG-NER complex in recruiting the Gcn5 onto chromatin in the correct regions of the genome to promote histone H3 acetylation required for the normal pattern of CPD repair rates observed in wild-type cells. This observation is striking because *GCN5* deleted cells are only moderately UV sensitive and exhibit near normal overall levels of repair of UV lesions. However, our experiments show that the genomic distribution of the rates of repair is markedly altered. We speculate that this could result in altered distributions of UV-induced genomic mutation patterns. If so, this could have important implications for genomic stability in tumourogenesis because cancer cells frequently display altered control of chromatin structure.

**Figure 5.**
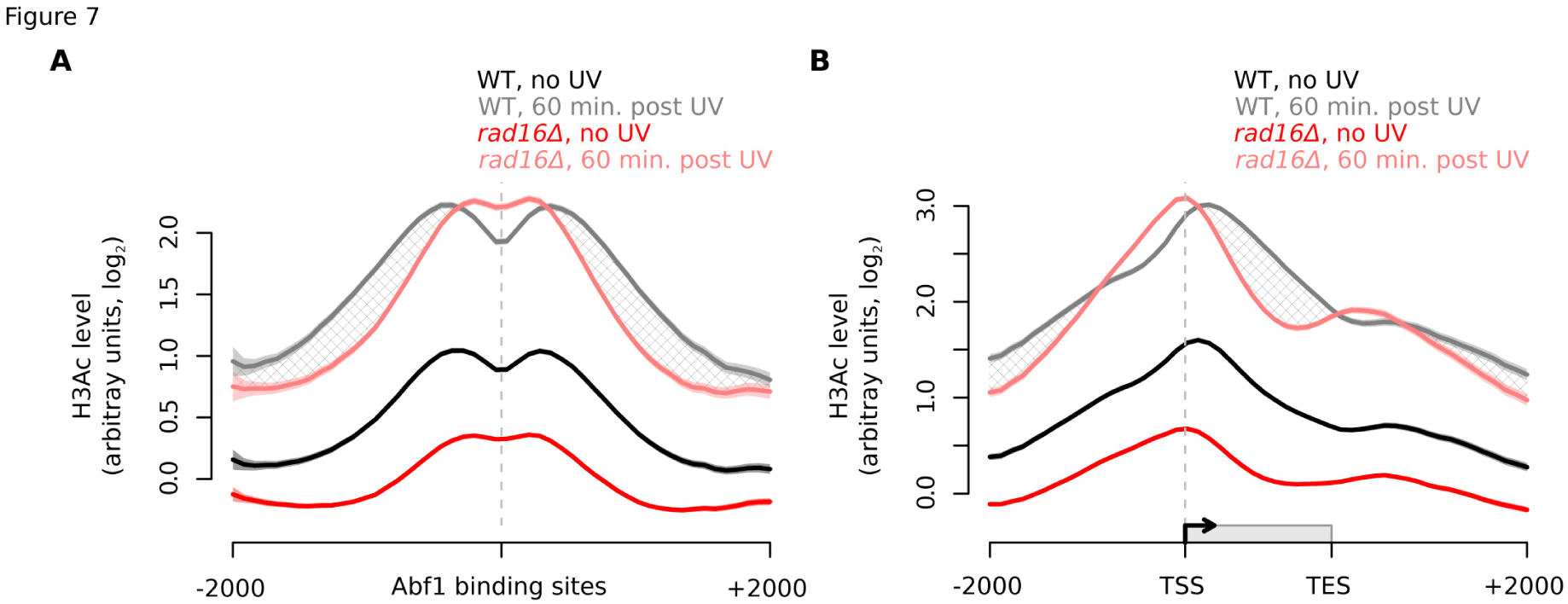
Histone H3 (K9, K14) acetylation levels in response to UV irradiation in wild-type and*rad16Δ* cells depend on the GG-NER complex. (**A**) Histone H3 acetylation in wild-type (n = 5, black/grey) and *rad16Δ* (n = 3, red/pink) cells in response to UV irradiation around ABF1 binding sites. The hatched areas define the genomic regions of GG-NER-dependent UV-induced histone H3 acetylation. Solid lines show the mean and shaded areas show the SEM. (**B**) As (A) plotted around ORF structure. The hatched areas define the genomic regions of GG-NER-dependent UV-induced histone H3 acetylation.

Recent reports have begun to measure and decipher the non-random nature of the mutational patterns that shape the somatic cancer genome that lead to disease. These include efforts to explain the causes of these mutation patterns based on our current knowledge of DNA damage and repair mechanisms (Haradhvala et al. 2016). Most recently, genomic DNA repair rates have been correlated with the incidence of mutations in skin cancer suggesting that cancer associated mutations occur in regions of the genome that are more difficult to repair (Adar et al. 2016; Perera et al. 2016; Sabarinathan et al. 2016). Our study demonstrates the importance of understanding the genomic organisation of DNA repair mechanisms in order to help explain the cellular processes that shape the cancer genome.

## Methods

### Strains and plasmids

| Plasmid/strain | Genotype | Reference |
| --- | --- | --- |
| *BY4742 (WT)* | MATα *his3Δ1 leu2Δ0 lys2Δ0 ura3Δ0* | EuroScarf |
| *rad16Δ* | *BY4742 rad16Δ::kanMX4* | EuroScarf |
| *rad7Δ* | *BY4742 rad7Δ::kanMX4* | EuroScarf |
| *GCN5-myc* | *BY4742 GCN5::myc9-URA3* | This Lab |
| *rad16ΔGCN5-myc* | *BY4742 rad16Δ::kanMX4 GCN5::myc9-URA3* | This Lab |
| *RAD7-myc* | *BY4742RAD7::myc18* | This Lab |
| *rad16Δ RAD7-myc* | *BY4742 rad16Δ::kanMX4 RAD7::myc18* | This Lab |
| *gcn5Δ* | *BY4742 gcn5Δ::kanMX4* | This Lab |
| *RAD16* | W303 *RAD16::myc18-URA3* | This Lab |
| *K216A* | W303 *RAD16K216A::myc18-URA3* | This Lab |
| *C552AH554A* | W303 *RAD16C552A,H554A::myc18-URA3* | This Lab |
| *K216AC552AH554A* | W303 *RAD16K216A,C552A,H554A::myc18-URA3* | This Lab |

### UV irradiation, yeast cell culture and crosslinking

Yeast cells were grown and UV irradiated as described previously (Yu et al. 2011). After the indicated repair time in YPD, 100 mL of cells were treated with formaldehyde to a final concentration of 1% for between 10-40 mins at room temperature to crosslink protein-DNA complexes. Crosslinking was stopped with the addition of 5.5ml Glycine (2.5 M) to 100 mL cells. Cells were collected by centrifugation and resuspended in cold PBS. For Rad7 capture using ChIP a double crosslinking method is required using 2×10^9^ cells collected by centrifugation and washed once with cold PBS. Crosslinking was carried out in two steps; first protein-protein crosslinking was performed using DMA (Dimethyl adipmidate dihydrochloride). Cells were resuspended in 10 mL of 10 mM and 0.25 % DMSO in cold PBS and incubated at room temperature for 45 min with mixing. Secondly, the cells were collected by centrifugation and washed 3 times with PBS and resuspended in 100 mL YPD. Finally, DNA-protein crosslinking was performed by addition of formaldehyde to a final concentration of 1%. The cell suspension was incubated for 45 min at room temperature.

### Chromatin preparation

Chromatin extracts were prepared as described previously (Teng et al. 2011; Yu et al. 2011). Briefly, cells were washed two times in cold PBS, followed by a wash step with cold FA/SDS (50 mM HEPES KOH pH 7.5, 150 mM NaCl, 1 mM EDTA, 1% Triton X-100, 0.1% NaDeoxycholate, 0.1% SDS, 1 mM PMSF). The cells were collected by centrifugation and prepared for lysis by bead beating in FA/SDS (+PMSF) using 0.5 mL of glass beads. The whole cell extract was then sonicated with a Bioruptor (Diagenode) as described previously (Yu et al. 2011), after which the chromatin extra was collected by centrifugation.

### Chromatin Immunoprecipitation

ChIP was performed as described previously (Yu et al. 2011; Powell et al. 2015). In short, Pre-washed pan mouse or anti-rabbit IgG Dynabeads were incubated with the respective antibody for 30 min at 30°C (2.5 μg of mouse anti-Myc (9E11, Abcam) antibody, 2.5 μL of rabbit anti-acetyl histone H3 (at K9 and K14, Upstate Biotechnology) or 20 μL anti-Abf1 antibody (yC-20, #sc-6679, Santa Cruz Biotechnology). Dynabeads were collected, washed and resuspended in 50 μL of PBS-BSA (0.1%) per sample. 30 μL of 10x PBS-BSA (10 mg/mL) and 100ul of sonicated DNA were added to each sample containing the Dynabeads, and the final volume made up to 300 μL with PBS. Samples were incubated at 21°C for 3 hours at 1300 rpm in an Eppendorf thermomixer. Following incubation, samples underwent a series of washes after which DNA was eluted from the Dynabeads in 125 μL of pronase buffer (25 mM Tris pH 7.5, 5 mM EDTA, 0.5% SDS) at 65°C at 900 rpm for 30 min. Pronase was added to each sample and incubated at 65°C in a water bath overnight. 50 μL of chromatin were taken as input samples and were treated with pronase and incubated overnight as the IP samples. The IP and input samples were treated with 5ul of DNase-free RNase A (10 mg/mL) for 1 hour at 37°C prior to DNA purification using the PureLink Quick PCR Purification Kit (Invitrogen) and eluted with 50 mL elution buffer.

### DNA preparation and IP for CPD detection

DNA was prepared and sonicated as described previously (Teng et al. 2011). IP was conducted as described in previous section (Chromatin Immunopreciptation) with the exception of using a different antibody for CPD IP (2 μg per sample of Anti-Thymine Dimer clone KTM53, Kamiya Biomedical Company). Following IP all samples were processed the same way to microarray.

### Removal of CPDs prior to microarray preparation and real-time PCR

CPDs need to be removed from the UV treated samples prior to PCR amplification and microarray hybridisation. The PreCR DNA repair kit (New England Biolabs) removes many DNA damages including CPDs. 40 μL of IP and IN sample were processed for repair using the PreCR repair kit as per the manufacturer’s instructions. Following repair, DNA was retrieved using phenol-chloroform extraction and ethanol precipitation and resuspended in 10 μL.

### DNA preparation and micro array hybridization

Samples were prepared for microarray hybridisation as detailed in the Agilent Technologies Yeast ChIP on chip protocol (Agilent Technologies Yeast ChIP-on-chip Analysis Protocol, version 9.2) and described previously (James’ 3D-DIP-chip paper, Sample Preparation for yeast DNA microarray hybridisation’ and ‘Microarray washing, scanning and feature extraction’ sections). Briefly, DNA fragments are blunt ended and a common linker sequence is ligated to their ends for two steps of ligation-mediated PCR amplification. The DNA concentration was measured with the NanoDrop spectrophotometer and adjusted to 150 ng/μL with H_2_O. 10.5 μL of IP and IN samples were differentially labelled with Alexa Fluor 5 and 3 dyes respectively, using the BioPrime Total Genomic Labelling System (Invitrogen). Labelling efficiency was determined using the MicroArray Measurement Module on the NanoDrop ND-1000 Spectrophotometer. The IP and input samples were combined and 110 μl hybridisation mixture applied to each Agilent yeast microarray. Mixtures were hybridised for 24 hours at 65°C. After hybridisation the microarrays were washed twice and scanned as described in the Agilent Technologies Yeast ChIP on chip protocol. The scanned image was processed with Agilent Feature Extraction software which converts fluorescence intensities into numerical values for analysis. Analysis of the data was conducted using Sandcastle (Bennett et al. 2015) in R version 3.2.4, which uses log_2_ values of IP:input ratios.

### Data Normalisation

Data from each experiment were normalised using the ‘normalise’ function in Sandcastle. The full Sandcastle normalisation procedure was applied to the individual protein binding and H3Ac datasets. This performs quantile normalisation on each set of replicates (for each time point), and then shifts and scales all datasets (combining all time points) together. This allows comparisons to be made between the binding levels of the protein/H3Ac at the different time points. Note that this procedure transforms the data so that for each set of protein/H3Ac datasets the values are comparable between the different time points analysed, but are not comparable between the different proteins/H3Ac. Only the quantile normalisation step was applied to each set of replicates of the CPD datasets, because these data are not suitable for the full Sandcastle normalisation procedure. For this reason the values of the damage levels at the different time points are not directly comparable and so the calculated repair rates are plotted using arbitrary units. It is thus possible to determine areas of relative faster or slower repair, but not to determine how much faster or slower repair is in one area compared to another.

### Data Analysis

The composite plots shown in this paper were created using the ‘profilePlot’ function of Sandcastle. Plots around Abf1 binding sites were created using peaks detected in the no UV treatment Abf1 binding datasets using the ‘enrichmentDetection’ function. Plots over ORFs were created using data downloaded from the ensembl databases using the ‘loadAnnotation’ function. Full details of these procedures are described in the Sandcastle publication and associated documentation (Bennett et al. 2015).

### Data Access

Data described here was submitted to EBIs Annotare 2.0 database and can be accessed using accession number E-MTAB-4641.

## Acknowledgements

The work was supported by an MRC Career Establishment Grant to SHR and a CRUK project grant A12340. SHR is also a member of the Cancer Genetics Biological Research unit funded by the Welsh Government.

### Author Contributions

The authors S.Y. and K.E. contributed equally.

### Disclosure Declaration

The authors declare no conflict of interest.

